# Exploring the phycosphere of *Emiliania huxleyi*: from bloom dynamics to microbiome assembly experiments

**DOI:** 10.1101/2022.02.21.481256

**Authors:** Mariana Câmara dos Reis, Sarah Romac, Florence Le Gall, Dominique Marie, Miguel J. Frada, Gil Koplovitz, Thierry Cariou, Nicolas Henry, Colomban de Vargas, Christian Jeanthon

## Abstract

Coccolithophores have global ecological and biogeochemical significance as the most important calcifying marine phytoplankton group. The structure and selection of prokaryotic communities associated with the most abundant coccolithophore and bloom-forming species, *Emiliania huxleyi*, are still poorly known. In this study, we assessed the diversity of bacterial communities associated with an *E. huxleyi* bloom in the Celtic Sea, exposed axenic *E. huxleyi* cultures to prokaryotic communities derived from bloom and non-bloom conditions and followed the dynamics of their microbiome composition over one year. Bloom-associated prokaryotic communities were dominated by SAR11, Marine group II Euryarchaeota, Rhodobacterales and contained substantial proportions of known indicators of phytoplankton bloom demises such as Flavobacteriaceae and Pseudoalteromonadaceae. Taxonomic richness of replicated co-cultures resulting from natural communities with axenic *E. huxleyi* rapidly shifted and then stabilized over time, presumably by ecological selection favoring more beneficial populations. Recruited microbiomes from the environment were consistently dependent on the composition of the initial bacterioplankton community. Phycosphere-associated communities derived from the *E. huxleyi* bloom depth were highly similar to one another, suggesting deterministic processes, whereas cultures from non-bloom conditions show an effect of both deterministic processes and stochasticity. Overall, this work sheds new light on the importance of the initial inoculum composition in microbiome recruitment and elucidates the temporal dynamics of its composition and long-term stability.

## Introduction

In the surface ocean, marine phytoplankton generate nearly up to 50% of global primary production and at least half of this production is remineralized by marine heterotrophic bacteria (Falkowski, 1994; Field et al., 1998; Pomeroy et al., 2007). From an ecological perspective, interactions between these essential microbial groups are being increasingly recognized as a major force shaping microbial communities (Amin et al., 2015, Seymour et al., 2017). Phytoplankton-bacteria interactions are widespread in marine environments, in particular within the phycosphere, the region immediately surrounding individual phytoplankton cell (Bell & Mitchell, 1972; Smriga et al., 2016). This microscale region, analogous to the plant root rhizosphere, serves as the interface for phytoplankton-bacteria associations. Phytoplankton exudates fuel the activity of heterotrophic microorganisms, that in exchange can stimulate microalgal growth through the provision of growth hormones and vitamins (Amin et al., 2015, Croft et al., 2005), protection against pathogenic bacteria (Seyedsayamdost et al., 2014) and through the facilitation of iron uptake (Amin et al., 2009). Phytoplankton release broad chemical classes of metabolites (Cirri & Pohnert, 2019) which can influence the taxonomy of phycosphere-associated bacteria (Buchan et al., 2014; Fu et al., 2020; Shibl et al., 2020).

Recent studies addressing the processes involved in bacterial community assembly in the phycosphere showed the influence of deterministic factors such as the place/time of isolation (Ajani et al., 2018) and the host species (Behringer et al., 2018; Lawson et al., 2018; Kimbrel et al., 2019; Sörenson et al., 2019; Jackrel et al., 2020; Mönnich et al., 2020). However, a combination between deterministic and stochastic effects in the microbiome recruitment process was also suggested (Kimbrel et al. 2019; Stock et al., 2022). To date, bacterial community composition and selection processes that influence the assembly of phycosphere microbiomes are not well known in many phytoplankton, in part because of the micrometer scale at which they take place (Kimbrel et al., 2019; Mönnich et al., 2020).

To overcome this challenge, one strategy is to study the selection processes in natural phytoplankton blooms (Zhou et al., 2019), in meso/microcosms or in cultures (Ajani et al., 2018; Kimbrel et al., 2019; Sörenson et al., 2019; Fu et al., 2020; Mönnich et al., 2020), when algal cells are at high concentrations. *Emiliania huxleyi* is the most abundant and cosmopolitan coccolithophore species and is able to form massive annual blooms in temperate and subpolar oceans mostly during Spring (Tyrrell & Merico, 2004). *E. huxleyi* blooms are characterized by blue turquoise waters that can be observed from satellite images (Tyrrell & Merico, 2004). These blooms have a critical importance for carbon and sulfur cycles due to the ecological and biogeochemical roles of coccolithophores as primary producers, calcifiers, and main contributors to the emission of dimethylsulfoniopropionate (DMSP) to the atmosphere (Malin & Steinke, 2004; Rost et al., 2004). The potential role of viruses in bloom termination has been thoroughly investigated (*e.g*. Bratbak, Egge & Heldal, 1993; Vardi et al., 2012; Lehahn et al., 2014), but only few studies have targeted the microbial diversity associated with *E. huxleyi* in natural environments (Gonzalez et al., 2000; Zubkov et al., 2001) and cultures (Green et al., 2015; Orata et al., 2016; Rosana et al., 2016). The *Roseobacter*, SAR86 and SAR11 lineages were identified as the main bacterial groups in natural *E. huxleyi* blooms (Gonzalez et al., 2000; Zubkov et al., 2001). Meanwhile, microbiomes of *E. huxleyi* in cultures are highly dominated by *Marinobacter* (Câmara dos Reis, 2021) and by Rhodobacteraceae (Green et al., 2015; Barak-Gavish et al., 2018).

In this study, we followed the dynamics of the prokaryotic community associated with *E. huxleyi* along a natural bloom in the Celtic Sea and used natural bloom and non-bloom samples to investigate the microbiome selection by an axenic *E. huxleyi* culture. We hypothesized that recruited microbiomes would be enriched in *Marinobacter* and Rhodobacteraceae members often associated with *E. huxleyi* cultures and would differ according to the initial prokaryotic composition.

## Materials and Methods

### Study site and sample collection

Samples used in this study were collected aboard the schooner *Tara* (Sunagawa et al., 2020) in the Celtic Sea (from 48°19-48°24 N/6°28-7°02 W; Fig. 1A and B), during the *‘Tara* BreizhBloom’ cruise from May 27 to June 2, 2019. To follow an *E. huxleyi* bloom formed in this area, an Argo float (https://argo.ucsd.edu/) was deployed in the center of the bloom patch and its position was used twice a day (early morning and end of the afternoon) to determine the geographical locations of the sampling stations.

**Fig. 1.**
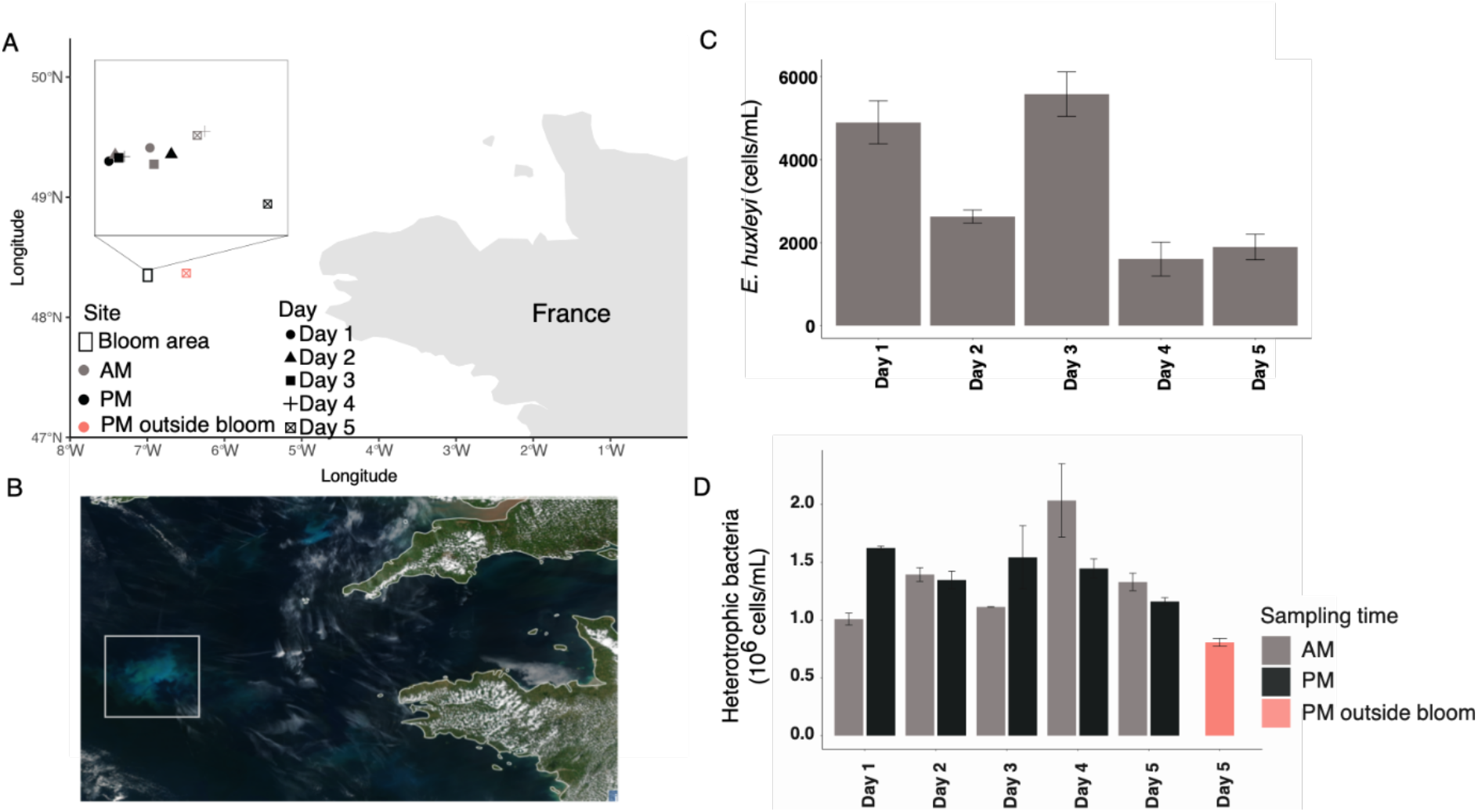
Sampling area and characteristics of the *E. huxleyi* bloom in the Celtic Sea. **A)** Map showing the bloom area and spatio-temporal sampling strategy. **B)** True-color satellite image of the bloom area on May 21, 2019 (source: https://www.star.nesdis.noaa.gov/sod/mecb/color/ocview/ocview.html. **C)** *Emiliania huxleyi* cell concentrations at morning bloom sites during the survey, measured from duplicate filters using scanning electron microscopy. **D)** Heterotrophic bacterial cell concentrations at morning and afternoon sites during the survey, measured from duplicate samples by flow cytometry.

On the last sampling day, a site about 34 km apart from the bloom area was also sampled (Fig. 1A). For each sampling event, surface to 50 m depth profiles of temperature, salinity, turbidity, pressure, photosynthetic active radiation (PAR), chlorophyll *a* (chla) fluorescence, oxygen concentrations and pH were conducted by deploying a SBE19+ profiler (Sea-Bird Scientific). Water samples were collected using an 8L Niskin bottle for nutrient analyses. After collection, nutrient samples (125 mL) were stored at −20°C for further analysis. Concentrations of nitrate, nitrite, phosphate and silicate were measured using a AA3 auto-analyzer (Seal Analytical) following the methods described by Tréguer & Le Corre, (1975) and Aminot & Kerouel (2007). Samples for flow cytometry (FCM), scanning electron microscopy (SEM), and metabarcoding analysis were collected at the bloom depth and prefiltered through a 20 μm mesh to eliminate large microzooplankton. For FCM analysis of photosynthetic eukaryotes and prokaryotic communities, two replicates (1.5 mL) were fixed using glutaraldehyde (0.25% final concentration) and Poloxamer 10% (0.1% final concentration) and incubated for 15 min at 4°C before flash freezing in liquid nitrogen. For SEM analysis, samples of morning sites (two replicates of 250 mL) were gently filtered onto PC membranes (47 mm in diameter; 1.2 μm pore-size) (Millipore). Filters were placed onto Petri slides, dried at least 2h at 50°C, and finally stored at room temperature. For metabarcoding analysis, cell biomass was collected from ~ 14 L of seawater by successive filtration onto large (142 mm in diameter) 3 μm pore-size and then 0.2 μm pore-size polycarbonate membranes (Millipore). Filters were flash-frozen in liquid nitrogen and stored at −80°C for later DNA analyses.

### Scanning electron microscopy analysis

Representative filter portions were fixed in aluminum stubs and sputter coated with gold–palladium (20 nm) (Keuter et al., 2019). Quantitative assessment of *E. huxleyi* cells was performed using a Phenom Pro scanning electron microscope. Cells were counted in twenty random screens (area analyzed = 0.16 mm^2^) and cell concentrations were calculated based on the filtered sample volume corresponding to the area analyzed (0.042 mL).

### Community assembly experiments

#### (i) Axenization

The *E. huxleyi* strain RCC1212, obtained from the Roscoff Culture Collection, was axenized following a sequence of washing and centrifugation steps, and variable incubation periods with increasing concentrations of an antibiotic solution mixture (ASM) as detailed in the original protocol developed at the Scottish Association for Marine Science (Oban, UK) available at: https://www.ccap.ac.uk/wp-content/uploads/2020/06/KB_Antibiotic_treatment.pdf. This method is briefly detailed in the Supplementary Materials and Methods section.

#### (ii) Sample preparation and inoculation

Four seawater samples were used in the bacterial community assembly experiment. They consisted of a surface and a DCM sample collected in the bloom area on day 5 (thereafter named inside bloom surface and inside bloom DCM) and a surface and a DCM sample collected the same day outside the bloom area (thereafter named outside bloom surface and outside bloom DCM) (Supplementary Fig. 1). In order to remove autotrophic picoeukaryotes and cyanobacteria from the inoculum, seawater samples were gently filtered through a 0.45 μm pore size membrane (Millex-HV, PVFD, Millipore). To estimate the number of prokaryotic cells lost during the filtration step, aliquots of total and filtered seawater samples were fixed for FCM analysis using the methods described in the Supplementary Materials and Methods. After filtration, 150 μL of each prokaryotic community were transferred in triplicates into 50 mL culture flasks filled with 15 mL of K/2 medium prepared as described in the Supplementary Materials and Methods. Finally, 150 μL of the axenic RCC1212 culture were added to each flask. Six flasks filled with 15 mL of K/2 medium and inoculated with 150 μL of the axenic RCC1212 culture were used as controls. In total, 18 cultures (12 treatments and 6 controls) were incubated at 15°C and a 12:12 photoperiod regime. Due to space limitation, only one thermostatic chamber with a light intensity of 20 μmol photons s^-1^m^-2^ using a blue neutral density filter was available onboard for incubation.

#### (iii) Survey of the culture microbiomes

Back to the laboratory and 10 days after inoculation, which corresponds to the time needed by *E. huxleyi* cultures to reach the end of exponential growth phase, axenic status of controls was checked by FCM. Then, each treatment was sampled for FCM analyses and prokaryotic community composition. Cultures (12 treatments and one axenic control) were transferred by inoculating 100 μL of the culture in 10 mL of fresh K/2 medium every 11-14 days for the first 176 days of experiment and then every 3 weeks until its end (day 393). The axenic control was regularly checked to ensure the clean handling of the cultures. In addition, culture flasks were randomized daily in the incubator to minimize positional effect on growth.

For FCM, duplicate samples were fixed as previously described and analyses performed according to Marie et al., (1999) are detailed in the Supplementary Materials and Methods section. For community composition analysis, 2 mL of culture was centrifuged at 2,000 *g* for 30 sec to reduce the microalgal load. Preliminary tests showed that this procedure reduces microalgal load while keeping most of the bacterial cells (about 90%). The supernatants were transferred into new tubes containing 2 μL of Poloxamer 188 solution 10% (Sigma-Aldrich) and centrifuged at 5,600 *g* for 5 min. The supernatants were discarded, and the pellets were stored at −80°C until DNA extraction.

### DNA extraction, PCR amplification and sequencing

DNA extraction from environmental and culture samples and amplification steps used to amplify the prokaryotic 16S rRNA gene using the universal prokaryote primers 515F-Y 5’-GTGYCAGCMGCCGCGGTAA-3’ (eight-nucleotide tag unique to each sample) and 926R 5’-CCGYCAATTYMTTTRAGTTT-3’ (Parada et al., 2016) are detailed in the Supplementary Materials and Methods and in (Romac, 2022a, 2022b, 2022c, 2022d). Library preparation and high-throughput sequencing using Illumina technology were performed at Fasteris SA (Plan-les-Ouates, Switzerland). Pooled amplicons were sent to Fasteris SA (Plan-les-Ouates, Switzerland) for Illumina high throughput sequencing.

### Bioinformatics

The steps of library separation, removal of Illumina adapters and first quality control were performed by Fasteris SA (see Supplementary Materials and Methods for details). The detailed scripts in this study (from bioinformatic treatment to statistical analysis) can be downloaded from https://github.com/mcamarareis/microbiome_assembly. Briefly, raw reads from each sequencing run were demultiplexed based on the 8 nucleotide tag sequences with cutadapt (version 2.8.1) (Martin, 2011). To deal with the presence of reads in mixed orientation in the R1 and R2 raw files (corresponding to each cycle of paired-end sequencing), the demultiplexing was performed in two rounds (see details in Supplementary Materials and Methods). Then, primer sequences were removed using cutadapt (version 2.8.1) (Martin, 2011). The demultiplexed primer-free sequences from different sequencing runs and from different sequencing cycles were processed independently to obtain an amplicon sequence variant (ASV) table using the DADA2 pipeline (version 1.14.0 in R 3.6.1) (Callahan et al., 2016; R Core Team, 2017). The ASV tables obtained independently for R1 and R2 were merged. Then, all the independent processed datasets were merged in one sequence table for chimera removal, also performed in pooled mode. The parameters used at each DADA2 step are specified in the Supplementary Table 1.

Sequences shorter than 366 bp and longer than 377 bp were filtered out, and the remaining sequences were taxonomically assigned using IDTAXA (50% confidence threshold) with the Silva database v138 (Quast et al., 2013; Murali et al., 2018). Chloroplasts and mitochondrial sequences were removed. Sequences not classified at the domain level by IDTAXA were assigned to the best hit in Silva v138 by pairwise global alignment (usearch_global VSEARCH’s command) (Rognes et al., 2016). These sequences were removed if they could not be classified and/or were classified as chloroplasts or mitochondrial sequences by VSEARCH (at 80% identity threshold). The resultant ASV table was filtered to remove ASVs accounting for less than 0.001% of the total number of reads. Consistency of technical replicates was evaluated by procrustes analysis, which measures the similarity between two ordinations of the same objects, followed by a protest, which measures the significance of the correlation. For this, we used a comparative principal components analysis performed on the Hellinger transformed data. After consistency was confirmed (p < 0.001 and correlation = 0.99), independent technical replicates of each culture were merged by the sum of the number of reads of the ASVs present in the two replicates of the same culture. The abundance and prevalence filters applied removed about 68% of the total number of ASVs while keeping 99% of the number of reads. The final dataset (filtered ASV table used for further analysis) contained 107 samples (11 from the environment and 96 from cultures) for a total of 6,017,019 reads and 294 prokaryotic ASVs.

### Community composition and statistical analyses

#### (i) Environmental samples

All the analyses were conducted in R version 4.0.2 in Rstudio (1.1.442) and the plots were produced with ggplot2 (RStudio Team, 2016; Wickham, 2016; R Core Team, 2017). Taxonomy treemaps of environmental samples were produced at the genus level considering the best hits classified by VSEARCH. To facilitate visualization, low abundant genera (accounting to less than 3% of relative abundance at each sample) present at the raw community table were grouped. To compare environmental samples and cultures, mean alpha diversity indices (richness and Shannon index) were measured after rarefying the ASV table 100 times at the minimum number of reads (2,479) (Saary et al., 2017). Hierarchical cluster analysis (HCA) (method “ward.D2”) was used to identify differences in the free-living prokaryotic communities by sites using the Hellinger distance (Euclidean distance of the Hellinger-transformed matrix) (Legendre & Gallagher, 2001; Oksanen et al., 2015).

#### (ii) Experiment

To analyze alpha diversity dynamics of the cultures, rarefaction was performed at a reads depth of 5,957 reads using the same approach as for the environmental samples. For beta diversity analysis, the Jaccard dissimilarity was calculated on the rarefied table (Oksanen et al., 2015). Hellinger distance was calculated from the non rarefied table (Legendre & Gallagher, 2001; Oksanen et al., 2015). To test the influence of the treatments, replicates and time on the microbiome beta diversity, we performed a permutational analysis of variance (PERMANOVA) (Anderson, 2005). Before running the analysis, the functions *betadisper* and *anova*-like permutation test from the package vegan were used to identify significant deviations on the multivariate beta dispersion of the data for treatments, replicates, time and of the interaction between treatments and time (Oksanen et al., 2015). The effect of treatments and replicates (nested within treatments) was tested using the function *nested.npmanova* from the package BiodiversityR (Kindt & Coe, 2005). To test the effect of time and the interaction between treatments and time, we used the function *adonis* (Oksanen et al., 2015) including treatments, replicates, and time (number of days) as fixed variables in the model, with permutations restricted to the replicates level. HCA was done using the Hellinger distance (Oksanen et al., 2015). Taxonomy barplots were produced by showing the three most abundant genera (considering the best hits classified by VSEARCH), while the less abundant were merged as “others”. *IndVal* analyses were run with the rarefied table to identify indicative species of the three groups of treatments evidenced in the beta diversity analysis (inside and outside bloom DCM and both surface samples) using the package indicspecies v1.7.9 with 10,000 permutations (De Cáceres and Legendre, 2009). P-values were adjusted for multiple comparisons using the false discovery rate method (Oksanen et al., 2015).

## Results

### Physico-chemical parameters and bacterial community structure dynamics in the *E. huxleyi* bloom

Coccolithophore blooms occur seasonally from April to June in the Bay of Biscay along the continental shelf to the Celtic Sea (Holligan et al., 1983; Poulton et al., 2014; Perrot et al., 2018). Here, we followed and sampled an *E. huxleyi* bloom for a week from end of May to early June 2019 in the Celtic Sea (Fig. 1A and 1B) using near-real time interpolated images of non-algal suspended particulate matter (SPM) derived from MERIS and MODIS satellite reflectance data (Perrot et al., 2018; Gohin et al., 2019) as provided by Ifremer (http://marc.ifremer.fr/en).

During the 5-day sampling period, temperature and salinity ranged from 12.4°C to 15.4°C and from 35.4 to 35.5 PSU, respectively (Supplementary Table 2). Nutrient concentrations were low with NO_2_ + NO_3_ and PO_4_ ranging from the detection thresholds to 1.25 μmol and 0.05 to 0.2 μmol/L, respectively. These low values were typical of a bloom event where cells consume most of the nutrients. *E. huxleyi* whose cell densities ranged from 1.6 x 10^3^ to 5.6 x 10^3^ cells/mL within the DCM layer of bloom waters (Fig. 1C) dominated the total photosynthetic eukaryotic community (2.5 x 10^4^ cells/mL on average). In these samples, total numbers of heterotrophic bacteria varied from 8.1 x 10^5^ to 2.0 x 10^6^ cells/mL (Fig. 1D) whereas the lowest prokaryotic cell concentration was measured in the outside bloom sample (Fig. 1D and Supplementary Table 2).

Overall, the inside and outside bloom DCM samples displayed a prokaryotic richness of about 140 ± 29ASVs (mean ± SD, n=11) (Fig. 2A). Richness increased over the course of the bloom, reaching a maximum at day 4. The DCM samples collected on day 5 inside and outside the bloom for the community assembly experiments contained 148 and 133 ASVs, respectively. The Shannon diversity index displayed homogeneous values (mean 4.1 ± 0.2) across samples (Fig. 2B).

**Fig. 2.**
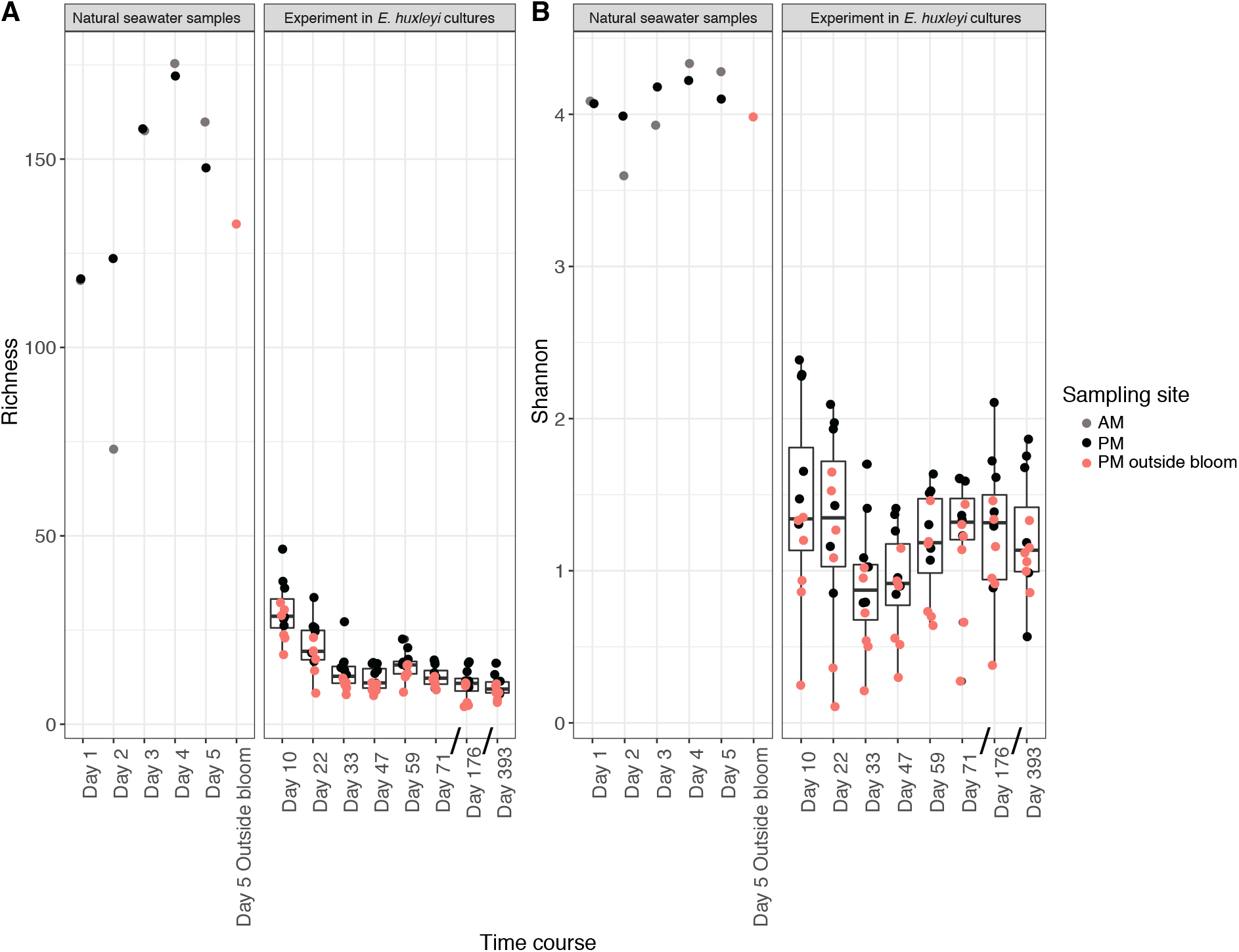
Composite representation of the dynamics of the prokaryotic richness **(A)** and Shannon **(B)** indexes in natural samples (0.2 - 3 μm DCM samples only) and culture experiments. ASV tables were rarefied to 2,479 (minimum number of reads of the environmental samples. The boxes represent the interquartile range. The thin horizontal lines represent the 25th and 75th percentiles and while the thick horizontal line represents the median. The vertical lines indicate the minimum and maximum values (using 1.5 coefficients above and below the percentiles). The dots represent the values measured for each culture. Dots further than the vertical lines represent potential outliers. X-axes in the in the experiment plot are not proportional to the time length between samplings.

Hierarchical clustering revealed 3 groups of samples that reflected the evolution of the prokaryotic communities across the bloom in a time-dependent manner (Supplementary Fig. 3). Free-living prokaryotic communities in and outside the bloom were mainly composed of members of Proteobacteria (66% of the total of reads), Bacteroidota (14%), Cyanobacteria (7%), Thermoplasmatota (4%), and Verrucomicrobiota (4%). Pelagibacteraceae (15%), Pseudoalteromonadaceae (13%) and Rhodobacteraceae (12%) were the most abundant proteobacterial families while Flavobacteriaceae (11%) dominated within the Bacteroidota. A diverse and rather stable prokaryotic community was observed at the genus level (Supplementary Fig. 4). Present in all samples, *Pseudoalteromonas* was the most abundant genus (13% of the total reads, n=11) and overdominated SAR11 inside (in several samples) and outside the *E. huxleyi* bloom. Abundances of SAR11 clade Ia (12% of the total reads), *Synechococcus* sp. (7%), SAR86 (5%), Marine group II Euryarchaeota (4%), uncultured Rhodobacteraceae members (4%), *Ascidiaceihabitans* (4%), and *Sulfitobacter* (4%) were also consistent along the study period. The Flavobacteriaceae dominated by the NS4 group (3%) comprised 11% of the total reads. Some genera dominated in only one sample like *Vibrio* (12% in day 2 PM sample) and *Nitrosopumilus* (18% in day 3 AM sample) or were substantial such as the members of OCS116 clade (4% in day 2 PM sample). Relative abundance of *Alteromonas* increased at the end of the bloom survey.

Prokaryotic communities outside the bloom grouped with those collected a few hours before inside the bloom area. The main compositional differences these samples that served as inocula for the community assembly experiment were the relative abundance of *Lentimonas* (5% inside vs 2% outside bloom), and of the OM60 (NOR5) clade and Marine group II members (each of them accounting for 2% inside vs 4% outside bloom).

### Community assembly experiment

#### (i) Dynamics of cell concentrations and alpha diversity patterns

Seawater samples used to inoculate axenic *E. huxleyi* cultures were filtered through 0.45 μm membranes to remove phototrophs (autotrophic eukaryotes and *Synechococcus* populations) and overcome their effects on microbiome assembly. FCM analysis demonstrated that about 40% of the initial bacterial cell concentration was lost after this filtration step. Due to limited incubation space onboard, cultures were incubated at low light (20 μmol photons s^-1^m^-2^) and these light conditions were maintained during the first weeks of incubation. However, a drastic decrease (~ 80%) of *E. huxleyi* cell concentrations was observed in all the treatments between the first (day 10) and the third (day 33) culture transfers (Supplementary Fig. 2A). To avoid culture crash, we increased the light intensity to 70 ± 20 μmol photons s^-1^m^-2^ and larger microalgal inocula (10% of the final culture volume instead of 1%) were used to transfer the cultures at day 33. *E. huxleyi* cell densities gradually increased at each subsequent transfer until day 71. At that point, they reached the highest cell concentration (9.9 x 10^5^ ± 1.2 x 10^5^ cells/mL) and remained stable up to the end of the experiment (Supplementary Fig. 2A). Bacterial cell concentrations followed an opposite trend during the first weeks of incubation. After a rapid increase (~94%) from day 10 to 47, they decreased once *E. huxleyi* abundance became higher and remained relatively stable up to the end of the experiment (Supplementary Fig. 2B).

Regarding the structure of the bacterial community, a severe loss of richness was observed between the environmental and culture samples (Fig. 2A). At day 10, the bacterial richness in the cultures was about one fifth of the richness in the natural samples (30 ± 8 SD, n=12) (Fig. 2A). This reflected a parallel decrease in the Shannon index, which at day 10, was about one third the values recorded in environmental samples (1.4 ± 0.6 SD, n=12, Fig. 2B). Over the course of the experiment, we observed a decrease in richness along the first five weeks (mean decrease 25 ± 7 SD, n=12, until day 47) (Supplementary Fig. 2C). After an increase at day 59 that corresponded to the period of culture recovery, the richness values decreased again and remained stable until day 393 (12 ASVs ± 2). The decrease of richness was mainly associated with the loss of low abundance ASVs, while the dominant ones remained over the course of the experiment (Supplementary Fig. 5). In general, the Shannon index also decreased over the first three time points (mean decrease 0.5 ± 0.6 SD, n=12) and then gradually increased to values similar to that from day 22 (day 393: 1.2 ±0.4, n=12). The highest richness and Shannon indexes were obtained in the treatments amended with the inside bloom DCM sample (richness: 26 ± 12 ASVs; Shannon 1.6 ± 0.4, n=24).

#### ii) Dynamics of beta diversity patterns in recruited microbiomes

In order to identify the influence of the different initial prokaryotic community composition and to follow the changes in the microbiome beta diversity with time, we used two metrics, the Hellinger distance (Euclidean distance of Hellinger-transformed data) and the Jaccard dissimilarity. Principal coordinates analysis using Jaccard dissimilarity demonstrated that *E. huxleyi* cultures inoculated with surface samples grouped together (Fig. 3A), while those inoculated with inside and outside bloom DCM samples formed two other independent clusters.

**Fig. 3.**
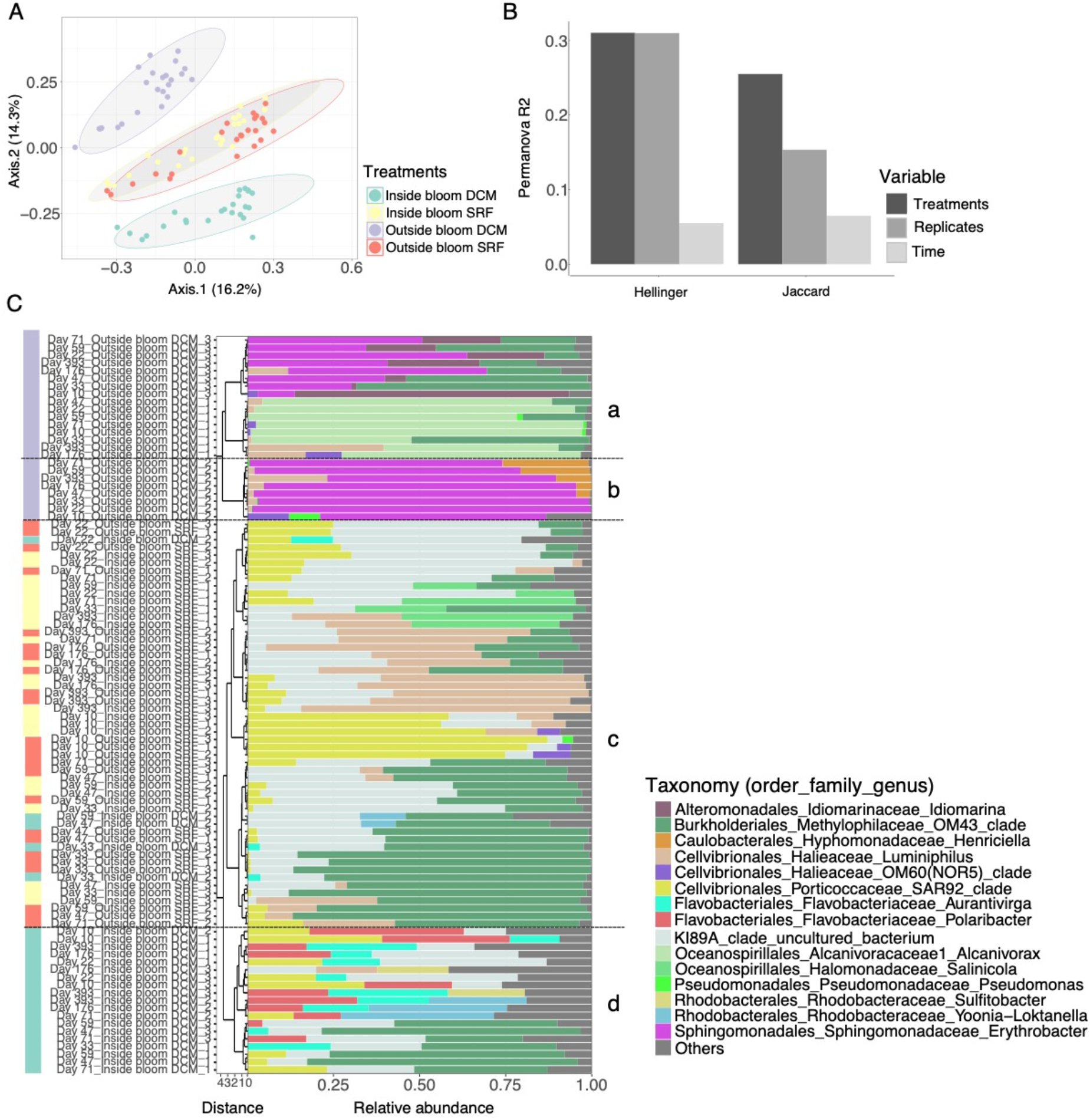
Beta diversity patterns of *E. huxleyi* microbiomes across treatments and time. (A) Principal coordinates analysis (PCoA) using Jaccard dissimilarity matrix of the presence-absence transformed rarefied table. Colors correspond to each treatment that received prokaryotic communities from different water samples: green - Inside bloom DCM; yellow - Inside bloom SRF; purple Outside bloom DCM; red - Outside bloom SRF. Ellipses represent 95% confidence. (B) r^2^ of permutational multivariate analysis of variance (PERMANOVA) and nested PERMANOVA using two metrics (see material and methods for details). (C) Hierarchical clustering produced with the Hellinger distance matrix using “ward.D2” method. Codes of each microbiome are experiment sampling day_treatment_replicate. Bar plots indicate the taxonomy of the 3 most abundant genera. The other genera were merged as “others”.

Statistical significance of the effect of treatments, replicates and time, as well as the interaction of treatment and time on the diversity of the microbiomes was assessed by PERMANOVA and nested PERMANOVA. Before performing PERMANOVA analysis we tested the beta-dispersion (variance) of the microbiomes grouped by treatments, time, treatments over time, and replicates. The dispersions (variance) of treatments as well as the interaction of treatments over time were not homogeneous for both metrics tested (p < 0.05). On the other hand, dispersions were likely homogeneous over time and across replicates (p > 0.05). Still, PERMANOVA results are robust to dispersion for balanced designs like ours (Anderson & Walsh, 2013). PERMANOVA results of Hellinger and Jaccard dissimilarities showed that significant proportions of the variance in microbiome composition among samples were explained by treatments (25-31%), replicates (15-31%), and time (6-7%) (p < 0.01) (Fig. 3B, Supplementary Tables 3 and 4). Although the interaction between treatments and time was significant using Hellinger distance (F = 2.23; p = 0.016), it explained a small proportion of the variance (2.5%).

Clustering using Hellinger distance revealed that the prokaryotic community composition of all the cultures grouped into four main clusters according to the inoculum origin (Fig. 3C). The outside bloom DCM treatment was formed by two clusters (a and b), highlighting the compositional differences between the 3 replicates. Replicate 1 (cluster a) was dominated by *Alcanivorax* (78%, n=8), while *Erythrobacter* prevailed in replicate 2 (cluster b, 88%) and 3 (cluster a, 45%). Cluster (c) mainly consisted of microbiomes recruited from both surface water samples. Surface treatments were dominated by bacteria related to OM43, KI89A, and SAR92 clades, and to *Luminiphilus*. The third main cluster (d) entirely formed by microbiomes from the inside bloom DCM treatment was dominated by members of the OM43 (29%), KI89A (28%), SAR92 clades, and *Polaribacter* (11%). ASVs that dominate (≥ 1%) in the cultures amended with inside and outside bloom DCM samples were generally low in abundance or not detected in the initial bacterioplankton community. However, relative abundance of cultured ASVs related to *Sulfitobacter*, *Luminiphilus* and the OM43 clade were also substantial in the field samples (Supplementary Fig. 6).

The number of indicative ASVs for each treatment varied widely and was significantly higher (21 out of 29) in cultures amended with inside bloom DCM waters (Supplementary Table 5). Flavobacteriales, with *Aurantivirga* and *Polaribacter* in particular, was the order containing the most indicative ASVs of microbiomes recruited from the inside bloom DCM sample. Members of SAR92 and KI89 clades displayed high Indval indexes in both inside bloom DCM and surface samples. The indicator ASVs of outside bloom DCM treatment were related to *Erythrobacter*, *Alcanivorax*, and OM60(NOR5) clade. Besides the compositional differences among treatments, we observed a somehow cyclic pattern of the beta-diversity over time using Hellinger distance (Fig. 4). Microbiome community compositions clearly differed from each other from days 10, 22 and 33 for all treatments, but they gradually tended to become similar to their initial status at the following time-points, particularly in the inside bloom DCM treatment (Fig. 4B). This pattern occurred at the replicate level in the outside bloom DCM cultures, and only for replicates 2 and 3 (Fig. 4D). This dynamic was mainly driven by the transient dominance of the OM43 clade (ASV1) during the alga crash and the increased abundance of *Luminiphilus* after the algal growth recovery (Supplementary Fig. 7).

**Fig. 4.**
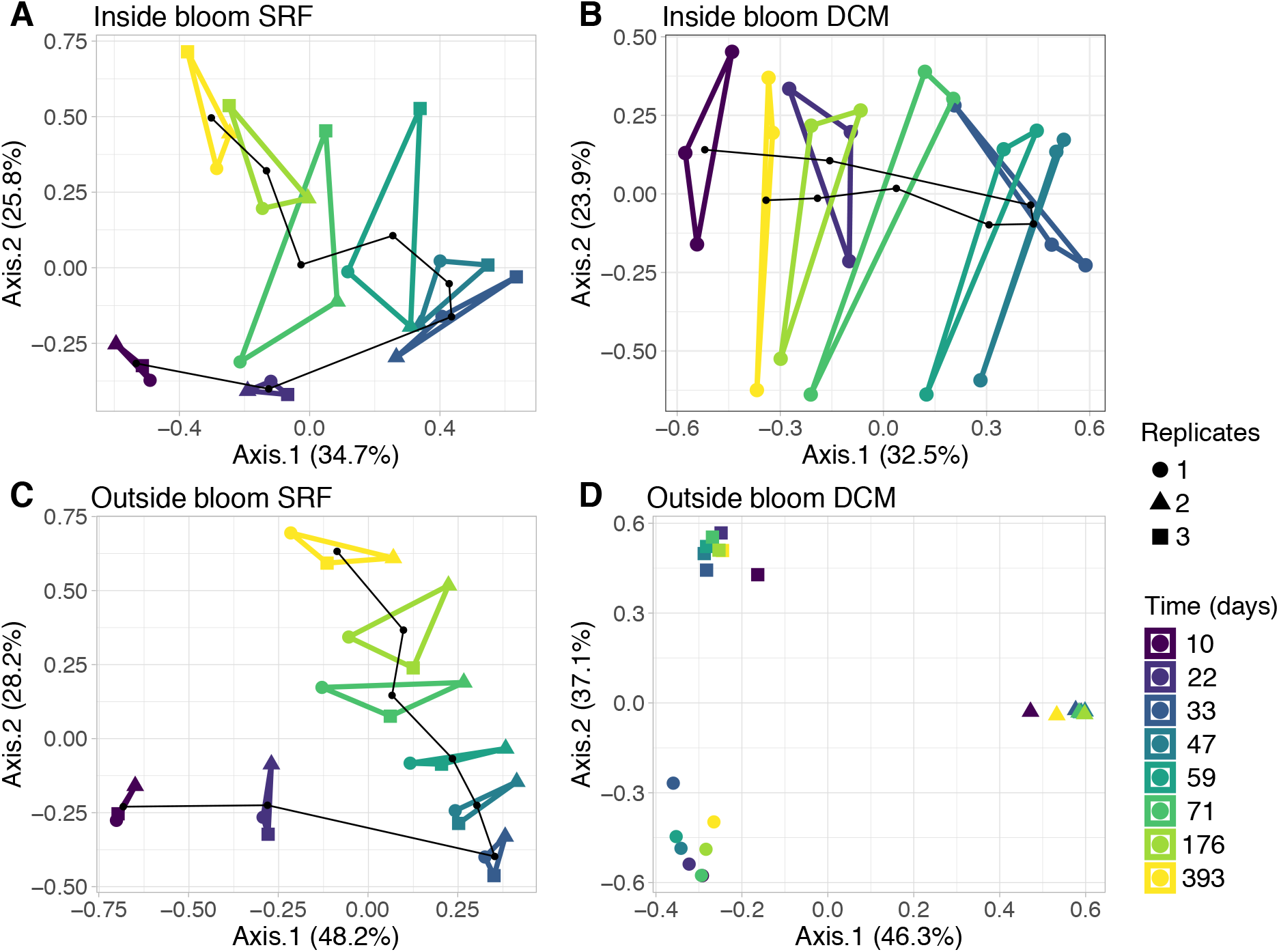
Principal components analysis (PCA) showing the cyclic patterns of the microbiome beta diversity. Community distances (Euclidean distances of Hellinger-transformed data) are shown for microbiomes from inside bloom SRF (A), inside bloom DCM (B), outside bloom SRF (C), and outside bloom DCM (D). The polygons link replicates (shape coded) at each time point (color coded). The black line links the barycenters of the replicates. In D., the cyclic pattern is observed at the replicate level for replicates 2 and 3.

## Discussion

In this study, we monitored the diversity of bacterial communities associated with an *E. huxleyi* bloom in the Celtic Sea, and collected bacterioplankton samples for conducting a microbiome selection experiment in axenic *E. huxleyi* cultures.

### Prokaryotic communities associated to the demise phase of the *E. huxleyi* bloom

Satellite images and post-cruise analyses indicate that the *E. huxleyi* bloom development had entered the decaying phase when we started the sampling. First, the high reflectance patch visible on the satellite images (Fig. 1A) and the daily vanishing of the coccolith-derived turbidity signal observed from the interpolated images of non-algal SPM were both indicative of detached coccoliths from dead *E. huxleyi* cells (Neukermans & Fournier, 2018; Perrot et al., 2018). This assumption was confirmed by the complete disappearance of the coccolith-derived turbidity signal a couple of days after we left the sampling area. Second, a suite of ongoing experiments on the bloom samples using diagnostic lipid- and gene-based molecular biomarkers (Vardi et al., 2009; Hunter et al., 2015; Ziv et al., 2016; Vincent et al., 2021) revealed the detection of specific viral polar lipids and visualized *E. huxleyi* infected cells during bloom succession, suggesting that Coccolithovirus infections may have partially participated in the demise of *E. huxleyi* bloom (F. Vincent, C. Kuhlisch, G. Schleyer, pers. comm.) as often proposed (Bratbak et al., 1993; Vardi et al., 2012; Laber et al., 2018). Third, the composition of the bacterial community, *i.e*. the presence of Flavobacteriaceae, Pseudoalteromonadaceae, Alteromonadaceae and members of the genus *Sulfitobacter*, is another indicator of bloom demise (Lovejoy et al., 1998; Buchan et al., 2014). Flavobacteriia, are reported amongst the main bacteria present in the declining phase of phytoplankton blooms (Teeling et al., 2012, 2016; Landa et al., 2016), which seems linked to their capacity to degrade high molecular weight substrates such as proteins and polysaccharides (Cottrell & Kirchman, 2000; Kirchman, 2002; Fernández-Gomez et al., 2013; Kappelmann et al., 2019; Francis et al., 2021). Finally, the algicidal effects of *Pseudoalteromonas, Alteromonas*, and *Sulfitobacter* strains and species have been documented in many microalgae including *E. huxleyi* (Holmström & Kjelleberg, 1999; Meyer et al., 2017; Li et al., 2018; Barak-Gavish et al., 2018), which calls attention to their potential role in the *E. huxleyi* bloom termination (Lovejoy et al., 1998; Barak-Gavish et al., 2018).

Our results evidenced a relative stability of the prokaryotic community during the study period and agree with reports of the co-occurrence of SAR11, *Roseobacter* and SAR86 clades in *E. huxleyi* blooms (Gonzalez et al., 2000; Zubkov et al., 2001). The co-occurrence of these groups could be mediated by the presence of dimethylsulfoniopropionate (DMSP), produced and released by *E. huxleyi* during blooms (Malin et al., 1993), which could be used as a sulfur compound by bacteria (Miller & Belas, 2004; Tripp et al., 2008; Dupont et al., 2012). DMSP and senescence compounds from decaying *E. huxleyi* cells probably also fueled members of the genus *Ascidiaceihabitans* (formerly Roseobacter OCT lineage) (Wemheuer et al., 2015), whose relative abundances typically fluctuate during phytoplankton blooms (Hahnke et al., 2015; Lucas et al., 2016; Chafee et al., 2018; Choi et al., 2018).

### Community composition of environmentally recruited *E. huxleyi* microbiomes

Since our primary objective was to study the bacterial community selection and assembly by a single phytoplankton host, we used a filtration step to discard autotrophic phytoplankton cells, such as *Synechococcus* and picoeukaryotes abundantly represented in the initial planktonic communities (Supplementary Table 2). We acknowledge that this filtration strategy has removed large and particle-attached prokaryotes, the latter probably being abundant in the demise phase of the bloom, and has induced substantial modifications in the initial community composition of the inocula and finally in recruited microbiomes. Indeed, some of the main taxonomic groups recruited in the treatments were not previously reported in *E. huxleyi* and other phytoplankton cultures or in low abundance, notably *Luminiphilus*, and the clades SAR92, KI89A and OM43 (Green et al., 2015; Câmara dos Reis, 2021). Except members of the OM43 clade (Yang et al., 2016), these bacteria are known as important groups of oligotrophic marine Proteobacteria that do not usually grow in the rich organic matter conditions provided in phytoplankton-derived cultures (Cho & Giovannoni, 2004; Spring & Riedel, 2013). Another unexpected result of our study is the very low representation of the genus *Marinobacter* in the recruited microbiomes whereas previous studies have reported their dominance in cultures of worldwide *E. huxleyi* isolates (Green et al., 2015; Câmara dos Reis, 2021). We cannot fully exclude the possibility that the filtration step impacted the abundance of *Marinobacter* in the inocula and finally in the *E. huxleyi* cultures. A more likely hypothesis however is that low light conditions might have induced algal cell death promoting the release of methylated compounds (Reese et al., 2019; Fisher et al., 2020). The release of methylated compounds by *E. huxleyi* may have provided a selective advantage to the specialist OM43 clade methylotrophs (Neufeld et al., 2008), dominant in all the cultures, and to other less common bacterial taxa in the absence of strong competitors such as *Marinobacter*. This hypothesis is in line with the opposite dynamics of *E. huxleyi* and bacteria coupled to the sharp decrease of the bacterial alpha diversity during the first month of culture, indicating that a few bacterial taxa were outcompeting others.

Despite the above limitations, the high reproducibility of microbiome community composition across the biological replicates suggests that they did not alter the general conclusions raised from our study.

Our results illustrate the importance of niche differentiation in natural communities. Indeed, although no major differences were observed between environmental inside and outside bloom bacterial communities, *E. huxleyi* microbiomes recruited from these samples differed. Similarly, although we did not analyze the initial bacterial composition of epipelagic surface samples (collected 34 km apart), they converged towards similar compositions, dominated by *Luminiphilus*, SAR92, KI89A and OM43 clades. Other microbiome studies of phytoplankton cultures have highlighted the impact of the initial community composition on microbiomes after short (Ajani et al., 2018; Sörenson et al., 2019; Jackrel et al., 2020), and long-term selection (Behringer et al., 2018). Remarkable features were found in the microbiomes resulting from inside bloom DCM waters where several indicative flavobacterial ASVs, mainly assigned to *Polaribacter* and *Aurantivirga*, were initially selected and remained among the most prevalent and abundant ASVs after growth recovery of the host. Both genera were identified as the main degraders of polysaccharides during diatom blooms (Krüger et al., 2019) and showed clear successions along the bloom stages (Teeling et al., 2012; Landa et al., 2016; Krüger et al., 2019; Liu et al., 2020). This may be related to the differential capacity of these bacteria to degrade phytoplankton-derived polysaccharides during blooms (Teeling et al., 2012; Krüger et al., 2019; Avci et al., 2020; Francis et al., 2021). Interestingly, SAR92 and *Luminiphilus* were also identified as important degraders of algal polysaccharides in bloom conditions (Francis et al., 2021), suggesting their potential functional role in our cultures.

We assume that exopolysaccharides/exudates of axenic *E. huxleyi* cultures have strongly influenced the initial microbiome composition and its long-term stability. This was exemplified by the contrasting results obtained with the inside and outside bloom DCM samples. The bacterioplankton communities associated with the inside bloom DCM sample displayed a higher diversity than that of the outside bloom DCM sample, and resulted in more diverse recruited microbiomes and higher number of indicative ASVs. We hypothesize that the bacterial assemblages adapted to *E. huxleyi* natural bloom conditions favored the recruitment of greater diversity microbiomes that may likely explain the almost complete cyclic pattern they followed (Fig. 4A). Such cyclic patterns were shown in the mucus microbiome of the coral *Porites astreoides* (Glasl et al., 2016) and the surface microbiome of the seaweed *Delisea pulchra* (Longford et al., 2019) after experimental disturbances. This pattern was consistently observed in the four separate treatments, with some variability. With the exception of the outside bloom DCM treatment, we observed lower levels of between-replicate variability at all time points, indicating a higher degree of uniformity among cultures at corresponding time-points.

Our study suggests the combined effect of deterministic processes and stochasticity on the microbiome assembly. The significant imprint of the original community in the inside bloom DCM treatments suggests that deterministic processes (*e.g*. assemblages adapted to *E. huxleyi* bloom exudates) influenced the final microbiome composition. On the other hand, variable communities selected from outside bloom DCM treatment show an effect of both deterministic processes and stochasticity. The latter microbiomes also contained commonly reported phytoplankton-associated bacterial groups (*Erythrobacter* and *Alcanivorax*), suggesting that the initial communities were exposed to other phytoplankton-mediated chemical conditions in the field. Variability in the community composition of biological replicates in this treatment suggests differential selection pressures, resulting in divergent communities. A similar pattern was found from enrichment experiments in the phycosphere of *Phaeodactylum tricornutum* (Kimbrel et al., 2019).

## Conclusions

In this work, we showed that the source of the initial bacterioplankton communities influences the resulting composition of *E. huxleyi* microbiomes. Our experimental approach demonstrated the stability of *E.huxleyi* microbiomes agreeing with previous reports for other phytoplankton microbiomes (Geng et al., 2016; Camp et al., 2020). Although species losses still occurred in the last sampled microbiomes of the survey, they were associated with low abundance taxa and did not induce major restructuring of the community, as previously shown in long-term experiments by Behringer et al., (2018). Overall, we bring evidence that microbiomes associated with phytoplankton cultures represent a valuable resource to explore phytoplankton-bacteria interactions. Isolation of indicative ASVs will be necessary to investigate the role and functions of stable core bacterial members interacting with *E. huxleyi*. Future co-culture experiments coupled with transcriptomic and metabolomic analyses will provide valuable information about the genes and molecules involved in these ecologically key interactions.

## Acknowledgments

This work was supported by a PhD fellowship from Sorbonne University and the Région Bretagne to MCdR, the Centre National de la Recherche Scientifique (CNRS, France), and the French Government ‘‘Investissements d’Avenir” programmes OCEANOMICS (ANR-11-BTBR-0008). We are grateful to the *Tara* Ocean Foundation, led by Romain Troublé and Etienne Bourgois, for the sampling opportunity and facilities onboard *Tara*, and to all the scientific and logistic team involved in the *Tara* Breizh Bloom cruise, notably captain Martin Herteau and his crew. We are thankful to the ABiMS bioinformatic platform (http://abims.sb-roscoff.fr) for the computational resources, to the RCC for providing the *E. huxleyi* cultures, and to Lydia White for her help in the statistical analysis.

## Data Accessibility

### Genetic data and sample metadata

Environmental samples are deposited in the bioproject PRJEB50692. Reads are deposited under the accession numbers XXXX and biosamples under the accession numbers ERS10466567-ERS10466596.

Samples from the assembly experiment are deposited in the bioproject PRJEB48747. Reads are deposited under the accession numbers XXXX and biosamples under the accession numbers ERS10539058-ERS10539153.

Unfortunately, we were unable to submit the reads of both bioprojects because of a problem on the ENA server. Sequences will be submitted once the server recovers.

## Author Contributions

CJ, MCdR, CdV designed the research. MCdR, CJ and SR participated on the sampling cruise. MCdR and CJ collected the DNA and transferred the cultures. FLG, SR, CJ produced the genetic data. MJF and GK analyzed the SEM filters. DM and MCdR analyzed the FCM samples. TC analyzed the nutrients samples. MCdR and NH analyzed the results. MCdR, CJ and CdV wrote the paper. All authors contributed to the discussions that led to the final manuscript, revised it and approved the final version.

