## Supplementary Material for "Exploring the phycosphere of *Emiliania huxleyi*: from bloom dynamics to microbiome assembly experiments"

**Table of Contents:**


| | |
|---|---|
| **Supplementary Materials and Methods** | Page 2 |
| **Supplementary Figure 1** | Page 7 |
| **Supplementary Table 1** | Page 8 |
| **Supplementary Table 2** | Page 8 |
| **Supplementary Figure 2** | Page 9 |
| **Supplementary Figure 3** | Page 10 |
| **Supplementary Figure 4** | Page 11 |
| **Supplementary Figure 5** | Page 12 |
| **Supplementary Table 3** | Page 13 |
| **Supplementary Table 4** | Page 13 |
| **Supplementary Figure 6** | Page 14 |
| **Supplementary Figure 7** | Page 15 |
| **Supplementary Table 5** | Page 16 |

### Supplementary Materials and Methods

#### Axenization:

Briefly, 15 mL of an exponential phase culture (~ $10^6$ cells/mL) were centrifuged at 1600 *g* for 4 min. The pellet was resuspended in sterile K/2 medium (Keller et al., 1987; Probert, 2019) prepared with reconstituted seawater (https://www.redseafish.com/red-sea-salts/red-sea-salt/; 37.7 grams of salts in 1L of ultrapure water, boiled 100°C for 20 min) and filtered through 0.1 µm. After centrifugation at 1600 *g* for 2 min, the supernatant was discarded, and the process was repeated for 6 times. The final pellet was resuspended in sterile K/2 medium and further used as inoculum. The axenization medium (2 mL of K/2 medium) contained increasing ASM concentrations (from 0 to 1X), and 25 µL of Marine Broth (1/10 strength) was added to promote bacterial growth. The 10X ASM contained the following antibiotics: cefotaxime (5 g/L), carbenicillin (5 g/L), kanamycin (2 g/L) and augmentin (2 g/L). The ASM was filtered (0.1 µm) and kept at -20°C for long-term storage. Culture tubes (one tube per ASM condition) inoculated with 100 µL of prewashed RCC1212 culture were incubated at 17°C with an irradiance of 70 ± 20 µmol photons $s^{-1}m^{-2}$. Aliquots (50 µL) of these cultures were sampled daily for 5 days and transferred into fresh K/2 medium and then incubated as before.

Once these new cultures were dense (about 15 days later), the presence of bacteria was checked by FCM. For FCM, 196 µL of each positive culture was fixed with glutaraldehyde 25% (0.25% final concentration) for 15 min in the dark. Then fixed cultures were stained with SYBR green (1/10,000 final concentration). Samples were analyzed using a FACSCanto flow cytometer (Becton Dickinson, San Jose, CA, USA) equipped with 488 and 633 nm lasers and standard filter setup. Bacterial data acquisition was triggered on the green fluorescence signal and non-diluted samples were run for 1 min at medium rate (~50 µL/min). Bacteria-free cultures were transferred into fresh K/2 medium and reinspected by FCM after the next culture cycle. For maintenance, axenic cultures were grown in K/2 medium prepared as previously mentioned. They were routinely checked for bacterial contamination by FCM as mentioned above. To ensure cultures remained axenic, culture aliquots (3-5 mL) stained with SYBR green and filtered onto 0.2 µm polycarbonate black membrane (Millipore Isopore) were examined by fluorescence microscopy.

**FCM analysis**

***(i) Environmental samples.*** Back to the laboratory, frozen samples were thawed at room temperature. Fluorescent microspheres (0.95 µm PolySciences) were added to each sample as internal reference at a final concentration between 6000 to 8000 per mL. Phytoplanktonic cells were first analyzed based on their autofluorescence using a FACSAria flow cytometer (Becton Dickinson, CA, USA) equipped with 488 and 633 nm lasers and the standard filter setup at a flow rate of 64 µL/min. A second analysis was run after staining samples with SYBR Green-I (1/10,000, final concentration) to enumerate heterotrophic microorganisms.

***(ii) Laboratory experimental samples.*** A FACSCanto flow cytometer (Becton Dickinson, San Jose, CA, USA) equipped with 488 and 633 nm lasers and standard filter setup was used to enumerate *E. huxleyi* and bacterial cells (Marie et al., 1999). For *E. huxleyi*, data acquisition was triggered on the red fluorescence signal and samples were run for 1 min at a flow rate of ~50 µL/min. To enumerate prokaryotes, samples were diluted 1:10 to 1:100 in TE (10 mM Tris, 1 mM EDTA [pH 8]), stained with SYBR Green (1/10,000 final concentration) and incubated for 15 min in the dark. The discriminator was set on green fluorescence, and the samples were analyzed as previously described.

**DNA extraction, PCR amplification and sequencing**

A mechanical cryogrinding step detailed in Romac (2022a) was applied to the 0.2 μm membranes collected onboard. DNA was extracted from the cryogrinded powders using the NucleoSpin RNA kit (Macherey-Nagel) combined with DNA elution buffer set using the protocol available in Romac (2022c). DNA from culture pellets was extracted using a modified protocol from NucleoSpin PlantII® DNA Mini kit (Macherey-Nagel) available in Romac (2022d). PCR (30 μL) was performed in triplicates using GoTaq G2 Flexi polymerase (Promega) following the method detailed in Romac (2022b). PCR products were purified using the NucleoSpin Gel and PCR Clean-Up kit (Macherey-Nagel), and quantified with the Quant-It PicoGreen double stranded DNA Assay kit (ThermoFisher). The purified PCR products were pooled in equal concentrations. In total three DNA pools were sequenced. The two first, containing environmental samples and the first 84 DNA samples (first 7 time points) were sequenced in two independent Illumina runs (technical replicates). The last 12 DNA samples (day 393) were sequenced in another Illumina run without sequencing replicates.

**Bioinformatics**

***(i)*** **Library separation, removal of Illumina adapters and first quality control.** First, in order to separate the libraries, the base calling was done based on a 6 nt unique index, using the softwares MiSeq Control Software 2.6.2.1, RTA 1.18.54 and bcl2fastq2.17 v2.17.1.14 and allowing 1 mismatch. Then, Illumina standard adapters removal and quality trimming were done using the Trimmomatic package (version 0.32) (Bolger et al., 2014). Briefly paired-end reads were globally aligned to ensure an end-to-end match. The adapter sequences were identified allowing a maximum of 2 mismatches and removed if the quality score was higher than 30. Then, bases were filtered by quality using a 4-base sliding window scan and trimming was done when the average quality per base dropped below 5. Reads without insert and with ambiguities were removed.

***(ii)*** **Correcting the mixed orientation of the reads.** Since R1 and R2 files (corresponding to each cycle of paired-end sequencing) contained reads with forward and reverse primers (further called forward and reverse reads), the demultiplexing was run two times. The first and second demultiplexing searched for adapters in the R1 file and in the R2 file, respectively, which separated forward and reverse reads for each sample corresponding to each cycle (Callahan et al., 2016; R Core Team, 2017). Since R1 and R2 cycles can have different error rates, they were analyzed independently during all the DADA2 processing. The same was done for the different sequencing runs. Following ASV inference, the forward and reverse reads of R1 were merged. To correct the mixed orientation, the reverse reads of the R2 were reverse complemented before merging with the forward reads.

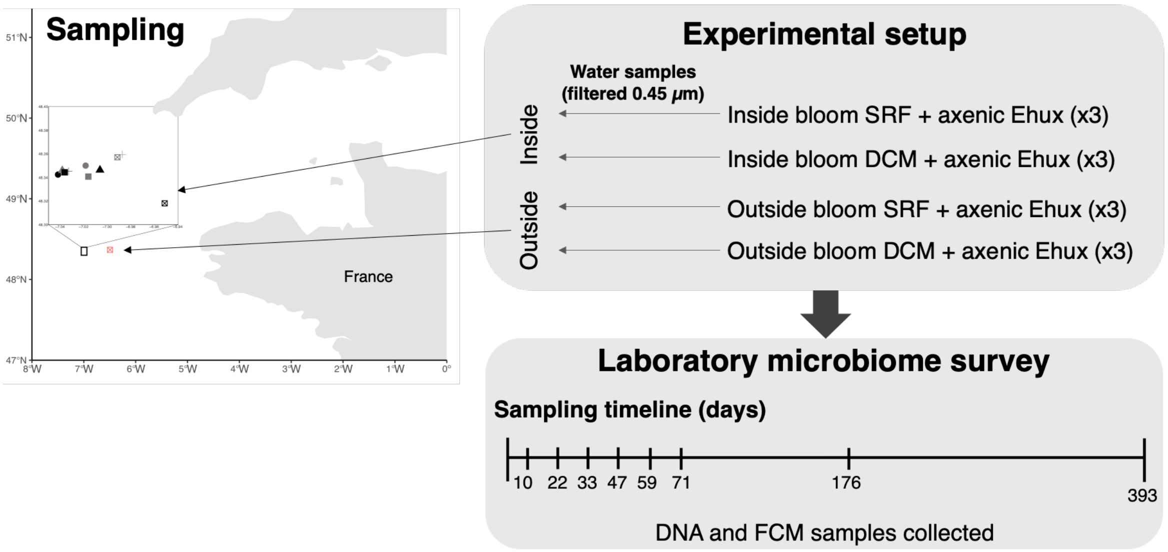


**Supplementary Figure 1.** Schematic view of sampling location, experimental design onboard, and molecular survey in the lab.

**Supplementary Table 1.** Settings used for each parameter in the DADA2 pipeline.

| | ***Reads from R1*** | ***Reads from R2*** |
|---|---|---|
| **filterAndTrim** | filterAndTrim(fastqFs1, filtFs1, fastqRs2, filtRs2, **truncLen=c(215,190)**, maxN=0, maxEE=c(2,2), truncQ=2, rm.phix=TRUE, compress=TRUE, multithread=TRUE) | filterAndTrim(fastqFs1, filtFs1, fastqRs2, filtRs2, **truncLen=c(190,215)**, maxN=0, maxEE=c(2,2), truncQ=2, rm.phix=TRUE, compress=TRUE, multithread=TRUE) |
| **learnErrors** | learnErrors(filtFs, nbases = 2e+08, randomize=TRUE, verbose=TRUE) / learnErrors(filtRs, nbases = 2e+08, randomize=TRUE, verbose=TRUE) | learnErrors(filtFs, nbases = 2e+08, randomize=TRUE) / learnErrors(filtRs, nbases = 2e+08, randomize=TRUE) |
| **dada** | dada(filtFs, err=errF, pool=TRUE) / dada(filtERs, err=errR, pool=TRUE) | dada(filtFs, err=errF, pool=TRUE) / dada(filtERs, err=errR, pool=TRUE) |
| **removeBimeraDenovo** | removeBimeraDenovo(seqtabs_merged, method="pooled",minFoldParentOverAbundance=8, verbose=TRUE, multithread=FALSE) | |

**Supplementary Table 2.** Environmental parameters and physico-chemical data of the cruise samples. The codes correspond to the day of sampling, the place where it was collected (inside - IN or outside - OUT of the bloom), the day time (morning - AM or afternoon - PM) and the depth (surface - SRF and deep chlorophyll maximum - DCM).

| Site code | *Date* | *Lat N* | *Long W* | *Time UTC* | *Sampling depth m* | *Nitrite µmol/L* | *Nitrate + nitrite µmol/L* | *Phosphate µmol/L* | *Silicate µmol/L* | *Temperature °C* | *Salinity PSU* | *Par* | *Fluorescence* | *Turbidity WETntu0* | *Oxygen Volts* | *Oxygen mL/L* | *Total Eukaryotes cells/mL* | *Synechococcus cells/mL* | *Heterotrophic bacteria cells/mL* |
|---|---|---|---|---|---|---|---|---|---|---|---|---|---|---|---|---|---|---|---|
| D1_IN_AM_DCM | 20190529 | 48°21.000 | 7°01.114 | 09:21 | 20 | 0.159 | 0.989 | 0.162 | 1.245 | 13.308 | 35.5042 | 6.55E+00 | 2.0457 | 1.8356 | 2.1988 | 5.3034 | 2.43E+04 | 4.20E+04 | 1.01E+06 |
| D1_IN_PM_DCM | 20190529 | 48°20.543 | 7°02.520 | 16:10 | 20 | 0.017 | DL | 0.078 | 0.986 | 14.2163 | 35.4004 | 2.01E+01 | 0.9833 | 2.0452 | 2.3406 | 5.56827 | 2.95E+04 | 6.40E+04 | 1.62E+06 |
| D2_IN_AM_DCM | 20190530 | 48°20.794 | 7°02.301 | 06:50 | 15 | 0.012 | 1.246 | 0.197 | 1.515 | 14.1133 | 35.3901 | 1.27E+01 | 0.7179 | 2.1609 | 2.1143 | 4.89932 | 2.46E+04 | 6.30E+04 | 1.39E+06 |
| D2_IN_PM_DCM | 20190530 | 48°20.817 | 7°00.384 | 16:31 | 20 | 0.021 | DL | 0.076 | 1.138 | 14.264 | 35.3993 | 3.42E+01 | 1.0677 | 2.0305 | 2.3299 | 5.52171 | 2.71E+04 | 6.25E+04 | 1.35E+06 |
| D3_IN_AM_DCM | 20190531 | 48°20.441 | 7°00.976 | 10:18 | 20 | 0.028 | 0.103 | 0.084 | 1.202 | 13.4539 | 35.376 | 5.85E+01 | 2.4072 | 1.9514 | 2.3633 | 5.73291 | 9.57E+03 | 2.08E+04 | 1.11E+06 |
| D3_IN_PM_DCM | 20190531 | 48°20.658 | 7°02.183 | 18:32 | 25 | 0.026 | DL | 0.068 | 1.117 | 13.2655 | 35.3729 | 1.44E+01 | 3.1488 | 2.048 | 2.335 | 5.6121 | 3.05E+04 | 7.48E+04 | 1.54E+06 |
| D4_IN_AM_DCM | 20190601 | 48°21.568 | 6°59.241 | 09:41 | 20 | 0.02 | DL | 0.083 | 1.067 | 12.4174 | 35.4077 | 1.51E+01 | 2.6518 | 1.8822 | 2.2428 | 5.39943 | 3.68E+04 | 1.10E+05 | 2.03E+06 |
| D4_IN_PM_DCM | 20190601 | 48°20.705 | 7°01.985 | 17:07 | 20 | 0.027 | 0.053 | 0.102 | 1.111 | ND | ND | ND | ND | ND | ND | ND | 3.21E+04 | 7.16E+04 | 1.45E+06 |
| D5_IN_AM_DCM | 20190602 | 48°21.427 | 6°59.496 | 09:40 | 15 | 0.035 | 0.04 | 0.061 | 0.922 | 13.2725 | 35.3667 | 8.69E+00 | 2.0143 | 1.9281 | 2.3174 | 5.58092 | 2.19E+04 | 1.01E+05 | 1.33E+06 |
| D5_IN_PM_SRF | 20190602 | 48°19.160 | 6°58.717 | 14:16 | 3 | 0.028 | 0.033 | 0.048 | 1.13 | ND | ND | ND | ND | ND | ND | ND | ND | ND | ND |
| D5_IN_PM_DCM | 20190602 | 48°19.081 | 6°57.090 | 13:45 | 25 | 0.03 | 0.035 | 0.094 | 0.276 | ND | ND | ND | ND | ND | ND | ND | 2.51E+04 | 9.15E+04 | 1.16E+06 |
| D5_OUT_PM_SRF | 20190602 | 48°21.961 | 6°28.519 | 18:00 | 3 | 0.034 | 0.039 | 0.073 | 0.197 | 15.3573 | 35.375 | 2.59E+02 | 0.4863 | 1.6543 | 2.3532 | 5.46434 | ND | ND | ND |
| D5_OUT_PM_DCM | 20190602 | 48°22.026 | 6°29.658 | 17:12 | 15 | 0.031 | 0.037 | 0.086 | 0.158 | 14.3971 | 35.3501 | 4.21E+01 | 1.3364 | 1.5285 | 2.3578 | 5.62631 | 1.05E+04 | 3.40E+04 | 8.06E+05 |

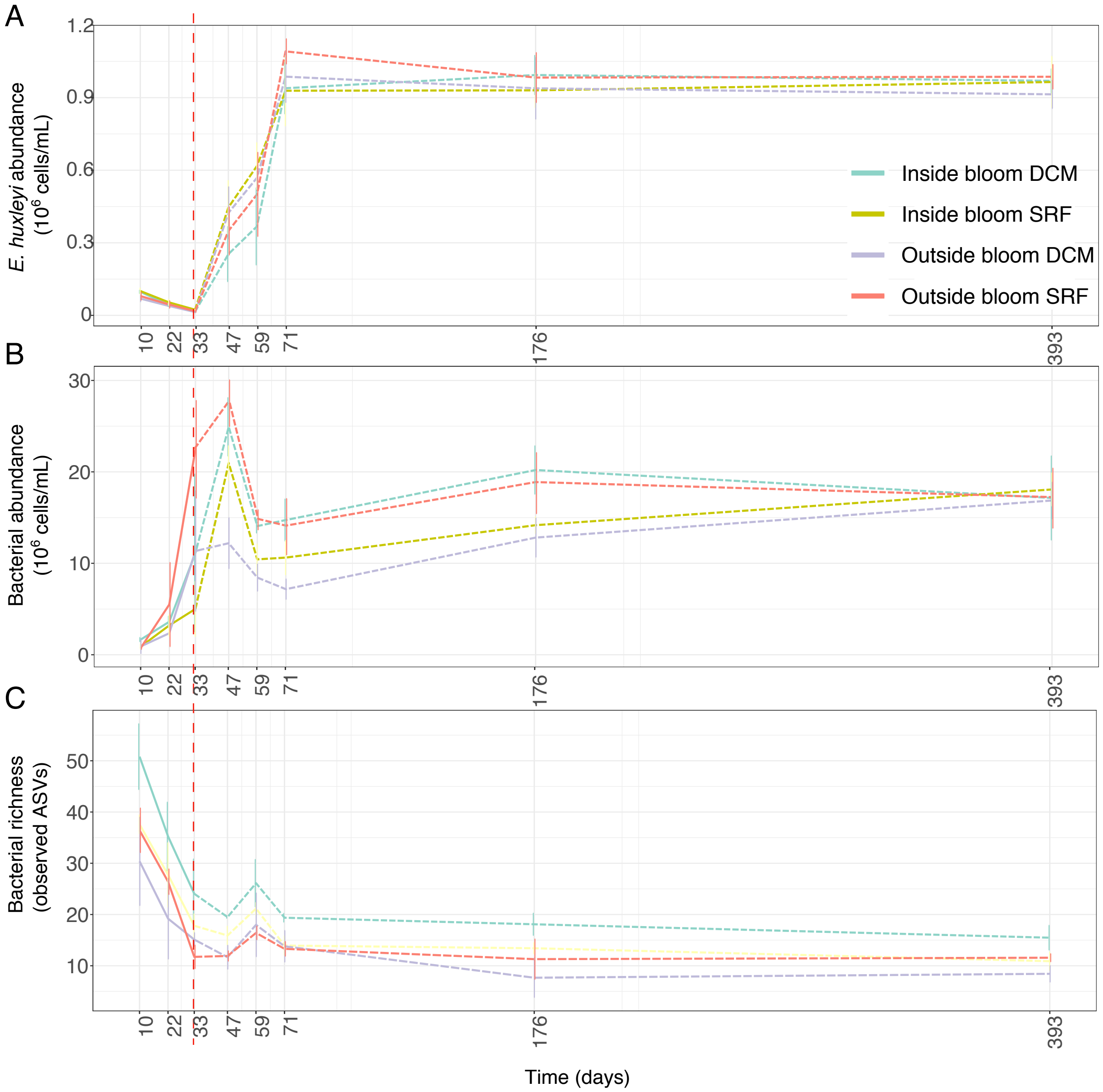


**Supplementary Figure 2.** Dynamics of **(A)** *E. huxleyi* and **(B)** bacterial cell concentration (cell/mL) over time (mean ± SD, n=3 for the four first time points and n=6 for the last four). **(C)** Richness, expressed as the number of prokaryotic ASVs over time for each treatment (mean ± SD, n=3). Colors correspond to each treatment that received prokaryotic communities from different water samples: green - Inside bloom DCM; yellow - Inside bloom surface; purple Outside bloom DCM; red - Outside bloom SRF. The red dotted line represents the moment following the increase of the inoculum at the culture transfer to recover *E. huxleyi* cell concentration.

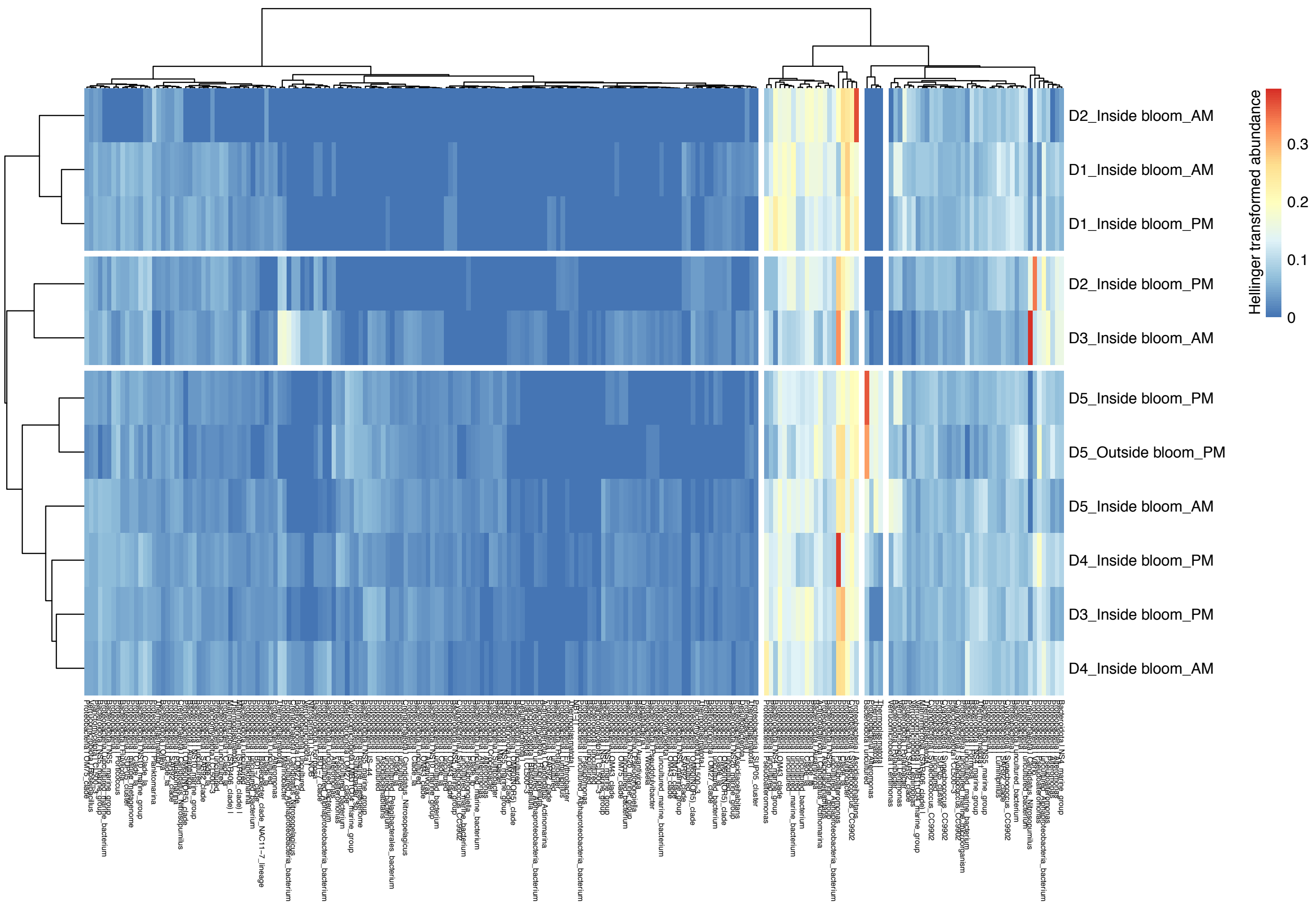


**Supplementary Figure 3.** Heatmap representing the hierarchical clustering of free-living (0.2-3 µm) prokaryotic communities at the DCM sampling sites. Samples codes refer to the day, depth, and time of sampling. The hierarchical clustering was built using the method "ward.D2" and Euclidean distances of the Hellinger-transformed community data. Taxonomy is represented by phylum and genus level. Taxonomic assignment at the genus level was performed usingVSEARCH (best hit). ASVs clustering is based in their relative abundance across samples.

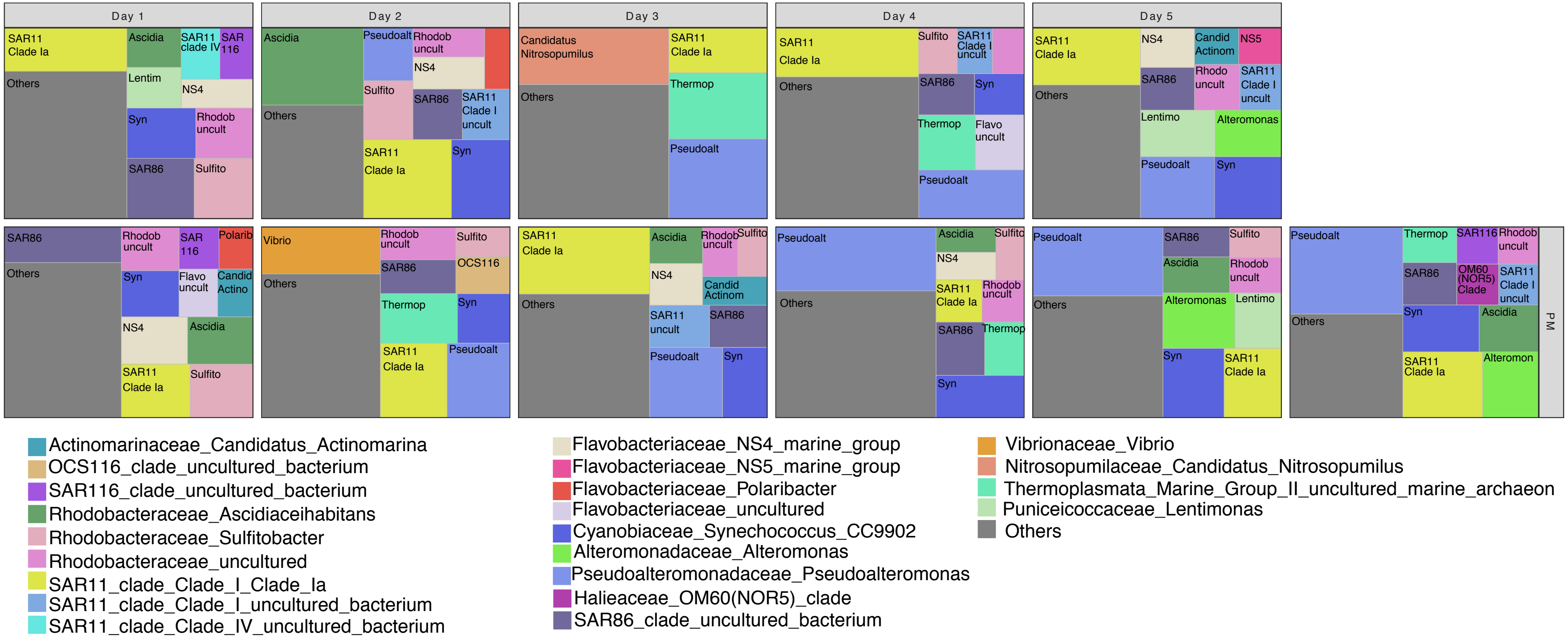


**Supplementary Figure 4.** Taxonomic composition of the free-living (0.2-3 µm) prokaryotic communities in DCM samples. Taxonomic assignment at the genus level was performed using VSEARCH (best hit). Genera accounting for less than 3% of the sample relative abundance were merged as “others”.

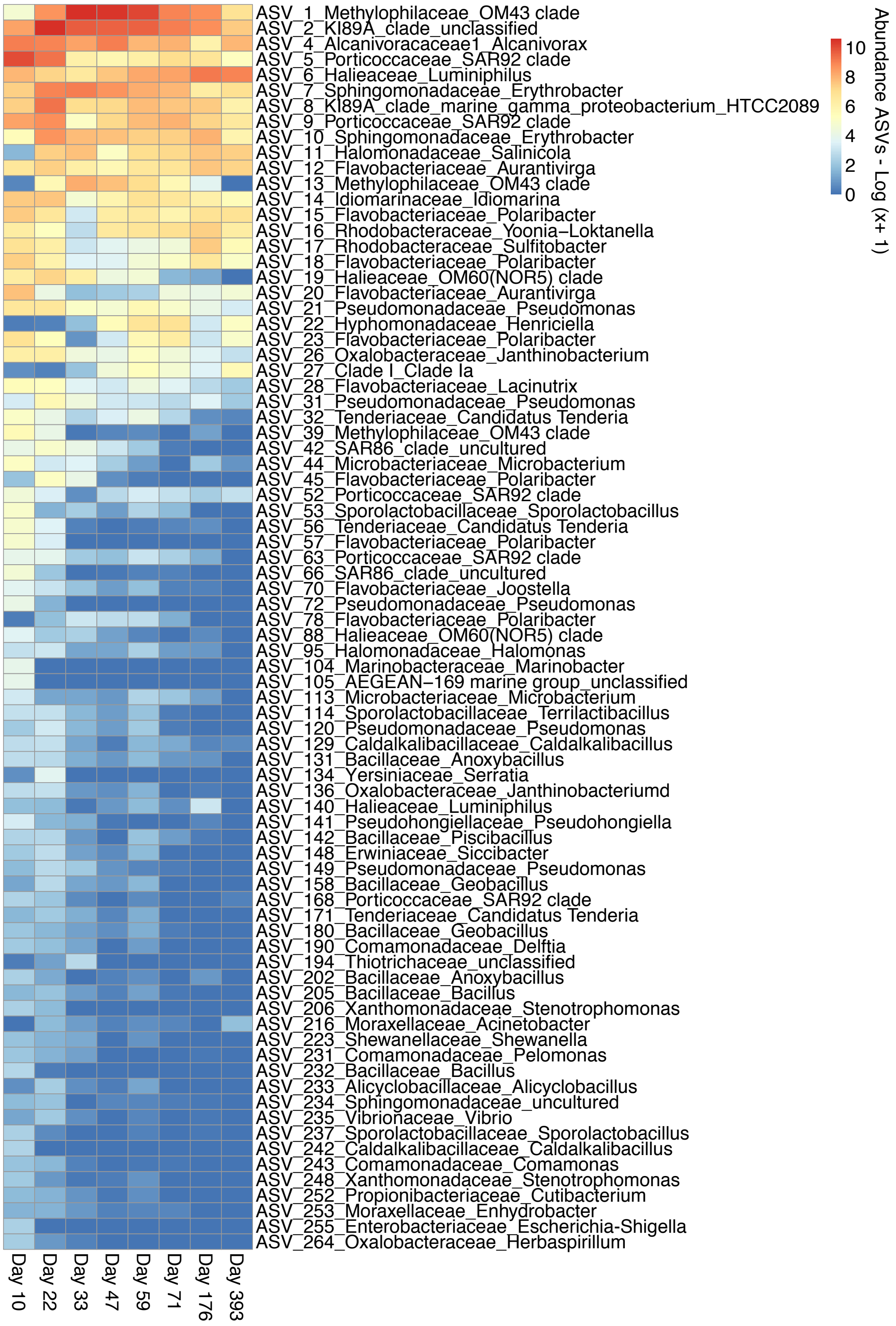


**Supplementary Figure 5.** Heatmap showing the ASV dynamics in bacterial consortia established in *E. huxleyi* cultures during the assembly experiment. ASV abundances correspond to the mean number of reads in all the samples at each time point. Abundances were log transformed - Log (x+1) - to facilitate visualization. Only ASVs with a mean number of reads higher than 10 are represented. Taxonomic assignment at the genus level was performed using VSEARCH (best hit).

**Supplementary Table 3.** Permutational multivariate analysis of variance (PERMANOVA) and nested PERMANOVA results (*nested.npmanova*) using Eulidean distance of Hellinger-transformed data. 'Treatments' is a fixed factor with four levels (inside bloom SRF, inside bloom DCM, outside bloom SRF and outside bloom DCM). 'Replicates' is a fixed factor with 12 levels (3 replicates x 4 treatments) nested within treatments. 'Time' is a fixed continuous factor (days after the starting of the experiment). Permutations were restricted to the replicates in traditional *adonis* (strata). •F.model and P-value correctly calculated using *nested.npmanova*. * Significant results.

| *Hellinger* | *Df* | *Sum of Squares* | *Mean sum of squares* | *F.Model* | *R2* | *P-value* |
|---|---|---|---|---|---|---|
| *Treatments•* | 3 | 17.944 | 5.981 | 2.670 | 0.310 | 0.001* |
| *Replicates•* | 8 | 17.923 | 2.240 | 8.561 | 0.310 | 0.001* |
| *Time* | 1 | 3.199 | 3.199 | 14.762 | 0.055 | 0.001* |
| *Treatments x Time* | 3 | 1.447 | 0.482 | 2.226 | 0.025 | 0.016* |
| *Residuals* | 80 | 17.337 | 0.217 | NA | 0.300 | NA |
| *Total* | 95 | 57.850 | NA | NA | 1 | NA |

**Supplementary Table 4.** Permutational multivariate analysis of variance (PERMANOVA) and nested PERMANOVA results (*nested.npmanova*) using Jaccard dissimilarity. 'Treatments' is a fixed factor with four levels (inside bloom SRF, inside bloom DCM, outside bloom SRF and outside bloom DCM). 'Replicates' is a fixed factor with 12 levels (3 replicates x 4 treatments) nested within treatments. 'Time' is a fixed continuous factor (days after the starting of the experiment). Permutations were restricted to the replicates in traditional *adonis*. •F.model and P-value correctly calculated using *nested.npmanova*. * Significant results.

| *Jaccard* | *Df* | *Sum of Squares* | *Mean sum of squares* | *F.Model* | *R2* | *P-value* |
|---|---|---|---|---|---|---|
| *Treatments•* | 3 | 5.904 | 1.968 | 4.402 | 0.254 | 0.001* |
| *Replicates•* | 8 | 3.576 | 0.447 | 2.730 | 0.154 | 0.001* |
| *Time* | 1 | 1.511 | 1.511 | 10.324 | 0.065 | 0.001* |
| *Treatments x Time* | 3 | 0.535 | 0.178 | 1.218 | 0.023 | 0.189 |
| *Residuals* | 80 | 11.709 | 0.146 | NA | 0.504 | NA |
| *Total* | 95 | 23.235 | NA | NA | 1 | NA |

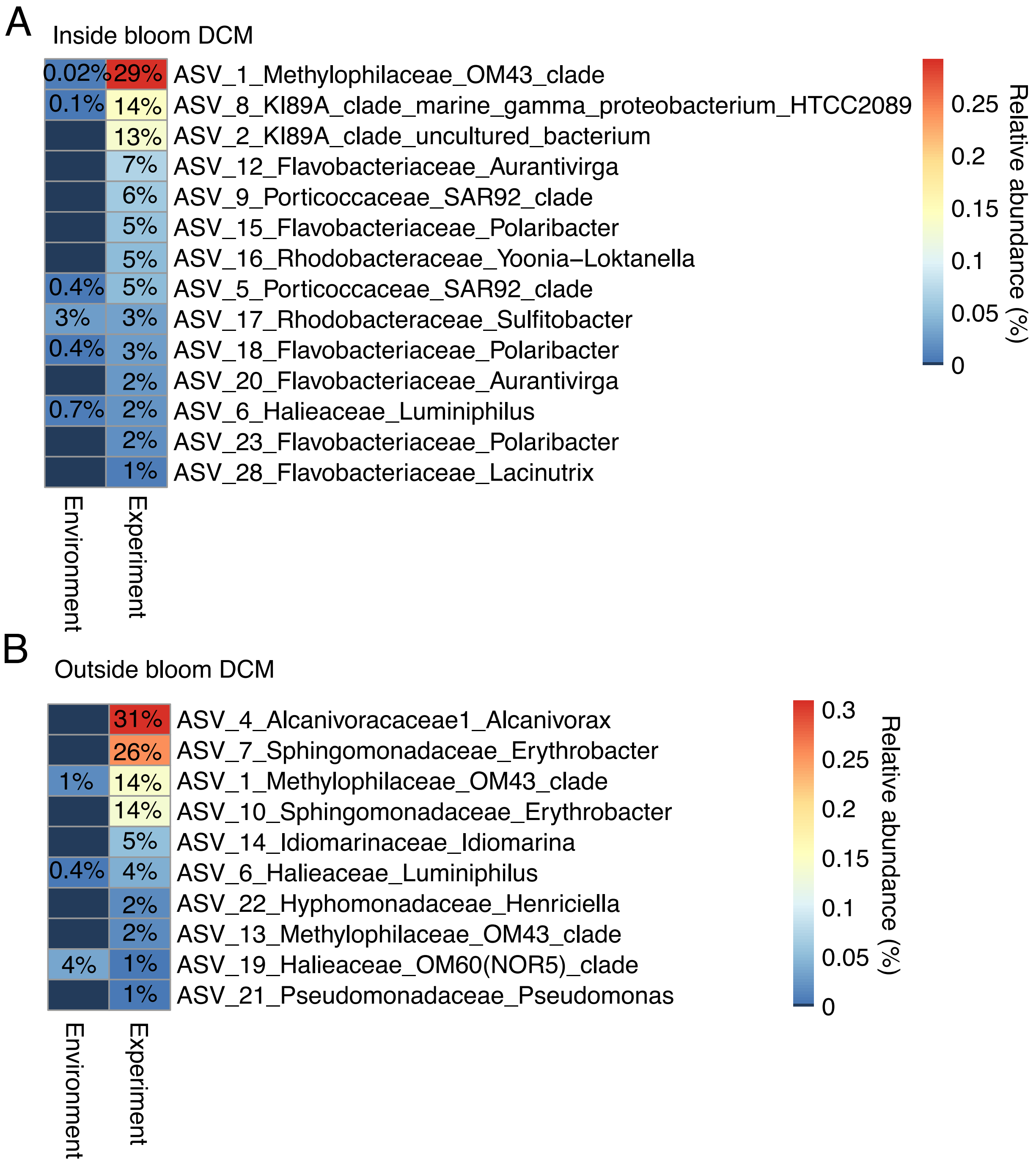


**Supplementary Figure 6.** Relative abundance of the dominant ASVs in cultures (experiment) and their corresponding proportions in the original samples (environment) of the inside bloom DCM (A) and outside bloom DCM (B).

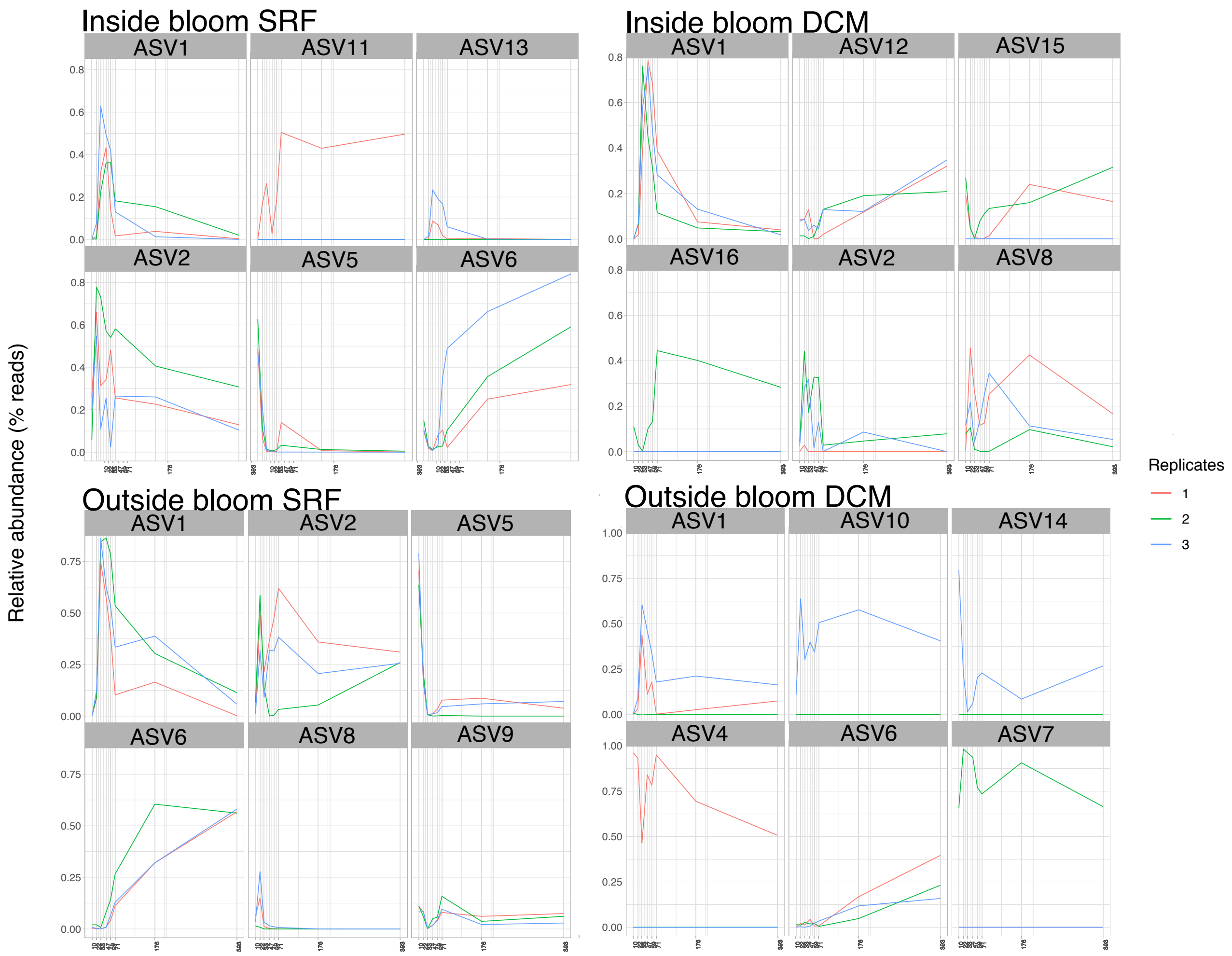


**Supplementary Figure 7.** Relative abundance dynamics of ASVs in the treatment triplicates over time (from day 10 to day 393). Relative abundance was calculated from the non-rarefied ASV table. ASV1 – OM43 clade, ASV2 – KI89A clade, ASV4 - *Alcanivorax*, ASV5 – SAR92 clade, ASV6 – *Luminiphilus*, ASV7 – *Erythrobacter*, ASV8 – KI89A clade, ASV9 – SAR92 clade, ASV10 – *Erythrobacter*, ASV11 – *Salinicola*, ASV12 – *Aurantivirga*, ASV13 – OM43 clade, ASV14 – *Idiomarina*, ASV15 – *Polaribacter*, ASV16 - *Yoonia-Loktanella*. Taxonomic assignment was performed using VSEARCH (best hit).

**Supplementary Table 5.** Detailed results of *IndVal* analysis. The grey-shaded polygons indicate the group to which the ASV is indicative. P-values were adjusted for multiple comparisons using false discovery rate correction.

| ASV | Taxonomy (order\|family\|genus) | Inside bloom DCM | Outside bloom DCM | SRF | *IndVal* | P-value | %Identity |
|---|---|---|---|---|---|---|---|
| ASV_12 | Flavobacteriales_Flavobacteriaceae_Aurantivirga | ■ | | | 1.000 | 0.0005 | 99.7 |
| ASV_18 | Flavobacteriales_Flavobacteriaceae_Polaribacter | ■ | | | 0.979 | 0.0005 | 100.0 |
| ASV_8 | KI89A_clade_marine_gamma_proteobacterium_HTCC2089 | ■ | | | 0.956 | 0.0005 | 100.0 |
| ASV_15 | Flavobacteriales_Flavobacteriaceae_Polaribacter | ■ | | | 0.913 | 0.0005 | 99.7 |
| ASV_20 | Flavobacteriales_Flavobacteriaceae_Aurantivirga | ■ | | | 0.890 | 0.0005 | 100.0 |
| ASV_23 | Flavobacteriales_Flavobacteriaceae_Polaribacter | ■ | | | 0.866 | 0.0005 | 99.7 |
| ASV_32 | Tenderiales_Tenderiaceae_Candidatus_Tenderia | ■ | | | 0.778 | 0.0005 | 100.0 |
| ASV_17 | Rhodobacterales_Rhodobacteraceae_Sulfitobacter | ■ | | | 0.677 | 0.0005 | 100.0 |
| ASV_16 | Rhodobacterales_Rhodobacteraceae_Yoonia-Loktanella | ■ | | | 0.577 | 0.0005 | 100.0 |
| ASV_28 | Flavobacteriales_Flavobacteriaceae_Lacinutrix | ■ | | | 0.577 | 0.0005 | 100.0 |
| ASV_58 | Flavobacteriales_Flavobacteriaceae_Polaribacter | ■ | | | 0.540 | 0.0005 | 100.0 |
| ASV_169 | Tenderiales_Tenderiaceae_Candidatus_Tenderia | ■ | | | 0.540 | 0.0005 | 97.1 |
| ASV_140 | Cellvibrionales_Halieaceae_Luminiphilus | ■ | | | 0.508 | 0.0146 | 99.7 |
| ASV_45 | Flavobacteriales_Flavobacteriaceae_Polaribacter | ■ | | | 0.500 | 0.0013 | 100.0 |
| ASV_78 | Flavobacteriales_Flavobacteriaceae_Polaribacter | ■ | | | 0.500 | 0.0013 | 99.7 |
| ASV_9 | Cellvibrionales_Porticoccaceae_SAR92_clade | ■ | | ■ | 1.000 | 0.0005 | 99.7 |
| ASV_2 | KI89A_clade_uncultured_bacterium | ■ | | ■ | 0.979 | 0.0005 | 99.7 |
| ASV_5 | Cellvibrionales_Porticoccaceae_SAR92_clade | ■ | | ■ | 0.972 | 0.0005 | 100.0 |
| ASV_64 | Cellvibrionales_Porticoccaceae_SAR92_clade | ■ | | ■ | 0.745 | 0.0010 | 100.0 |
| ASV_39 | Burkholderiales_Methylophilaceae_OM43_clade | ■ | | ■ | 0.707 | 0.0005 | 100.0 |
| ASV_206 | Bacillales_Bacillaceae_Bacillus | ■ | ■ | | 0.451 | 0.0282 | 100.0 |
| ASV_7 | Sphingomonadales_Sphingomonadaceae_Erythrobacter | | ■ | | 0.677 | 0.0005 | 100.0 |
| ASV_19 | Cellvibrionales_Halieaceae_OM60(NOR5)_clade | | ■ | | 0.629 | 0.0248 | 100.0 |
| ASV_4 | Oceanospirillales_Alcanivoracaceae1_Alcanivorax | | ■ | | 0.577 | 0.0005 | 100.0 |
| ASV_10 | Sphingomonadales_Sphingomonadaceae_Erythrobacter | | ■ | | 0.577 | 0.0005 | 100.0 |
| ASV_14 | Alteromonadales_Idiomarinaceae_Idiomarina | | ■ | | 0.577 | 0.0005 | 100.0 |
| ASV_22 | Caulobacterales_Hyphomonadaceae_Henriciella | | ■ | | 0.500 | 0.0017 | 99.2 |
| ASV_70 | Flavobacteriales_Flavobacteriaceae_Joostella | | ■ | | 0.500 | 0.0020 | 100.0 |
| ASV_13 | Burkholderiales_Methylophilaceae_OM43_clade | | ■ | ■ | 0.796 | 0.0321 | 100.0 |
